# Wide genome involvement in response to long-term selection for antibody response in an experimental population of White Leghorn chickens

**DOI:** 10.1101/046953

**Authors:** Mette Lillie, Zheya Sheng, Christa Honaker, Ben Dorshorst, Paul Siegel, Örjan Carlborg

## Abstract

Long-term selection experiments provide a powerful approach to gain empirical insights into adaptation. They allow researchers to uncover the targets of selection and how these contribute to the mode and tempo of adaptation. Here we report results from a pooled genome re-sequencing study to investigate the consequences of 39 generations of bidirectional selection in White Leghorn chickens on a humoral immune trait: antibody response to sheep red blood cells. We observed wide genome involvement in response to this selection regime, with over 200 candidate sweep regions characterised by spans of high genetic differentiation (*F*_*ST*_). These sweep signatures, encompassing almost 20% of the chicken genome (208.8 Mb), are primarily the result from bidirectional selection on haplotypes present in the base population. These extensive genomic changes highlight both the extent of standing genetic variation at immune loci available at the onset of selection, as well as how the long-term selection response results from selection on a highly polygenic genetic architecture. Furthermore, we present three examples of strong candidate genes that may have contributed to the profound phenotypic response to selection.

**Data Availability:** Pooled genome data generated for this study will become available via SRA upon acceptance of manuscript

## Introduction

A key aim in the study of evolutionary genetics is to identify loci that facilitate adaptation. The importance of de novo mutations and standing genetic variation to the mode and tempo of the adaptive process has been debated, as well as imporatnce of fixation for adaptation (Jensen 2014; Chevin and Hospital 2008; Pritchard et al. 2010). Through necessity, de novo mutations are essential within bacterial systems, where beneficial mutations arising within clonal populations will sweep to fixation. Interaction and competition between de novo mutations observed in experimental populations have provided valuable insights into the process of adaptation within microbial species (Burke 2012; Elena and Lenski 2003; Barrick et al. 2009). Research in sexual organisms, however, reveals that selection on standing genetic variation is the predominant basis of adaptation in higher eukaryotes (Burke et al. 2014; Hernandez et al. 2011; Hermisson and Pennings 2005; Sheng et al. 2015). In this case, standing genetic variants are present in the population at low frequency, maintained by neutral or slightly negative selection. Within an altered environment, these standing variants will gain a selective advantage and increase in frequency in the population to reach fixation. However, depending on the genetic architecture of functional traits and population structure, adaptation can also be facilitated by moderate allele frequency differences at multiple genes, without producing such dramatic sweep-to-fixation signatures (Pritchard and Di Rienzo 2010; Pritchard et al. 2010; Chevin and Hospital 2008).

Immune genes are known to evolve under balancing selection, a selective process where fixation is avoided and many haplotypes are maintained in populations (Nei et al. 1997; Hughes and Nei 1988). The first evidence for this came from studies of the major histocompatibility complex (MHC), where extremely high MHC diversity results from complex selective pressures exerted by diverse pathogen communities (Parham and Ohta 1996; Piertney and Oliver 2006; Hughes and Yeager 1998). This coevolutionary balancing selection also influences many genes involved in host defense responses, such as innate and adaptive immunity in vertebrates (Ferrer-Admetlla et al. 2008; Bamshad et al. 2002; Barreiro et al. 2009) and resistance genes in invertebrates and plants (Ghosh et al. 2012; Tian et al. 2002; Meyers et al. 2005). An important consequence of this balancing selection is that most populations are expected to maintain extensive standing immunogenetic variation, available for adaptation to new environmental and pathogenic challenges.

The number of genes underlying an adaptive process often belies the complexity of the selective environment. For traits with a complex genetic basis, such as aging and courtship, experimental evolution studies in *Drosophila* have demonstrated a genome-wide involvement in response to selection (Remolina et al. 2012; Turner et al. 2013). For immune traits, the extent of genome involvement in the adaptive response can vary. For example, in *Drosophila melanogaster*, nearly 5% of the genome, and 42 genes, were suggested to be involved in an 150% increase in parasitoid resistance (Jalvingh et al. 2014), whereas only few genes were identified and functionally validated as the targets of selection for resistance to *Drosophila* C virus (Martins et al. 2014).

Here we use a genomic approach to study the genome-wide consequences of long-term, bidirectional selection on a single immune trait from a base population of randombred White Leghorn chickens (Siegel and Gross 1980). In brief, selection was performed for high (HAS) or low (LAS) day 5 antibody production to an intravenous challenge of sheep red blood cells (SRBC) (further details can be found in Siegel and Gross 1980; Siegel et al. 1982; Boa-Amponsem et al. 1997). At generation 39, the HAS and LAS lines showed an average 6.5 fold difference in antibody titers (Figure 1). Pooled genome sequencing was carried out for each selected line at generation 39 (HAS39 and LAS39) allowing the identification of regions with high differentiation (*F*_*ST*_). Due to the bi-directional selection regime, the identified selection signatures would result primarily from selection on standing genetic variants (Johansson et al. 2010; Sheng et al. 2015). Relaxed lines (random-mated sublines) founded for both lines at generation 24 were also pool genome sequenced at generation 16 (HAR16 and LAR16). The genetic effects of the sweeps were also estimated using data from an F_2_ intercross between the HAS and LAS lines at generation 32.

**Figure 1.**
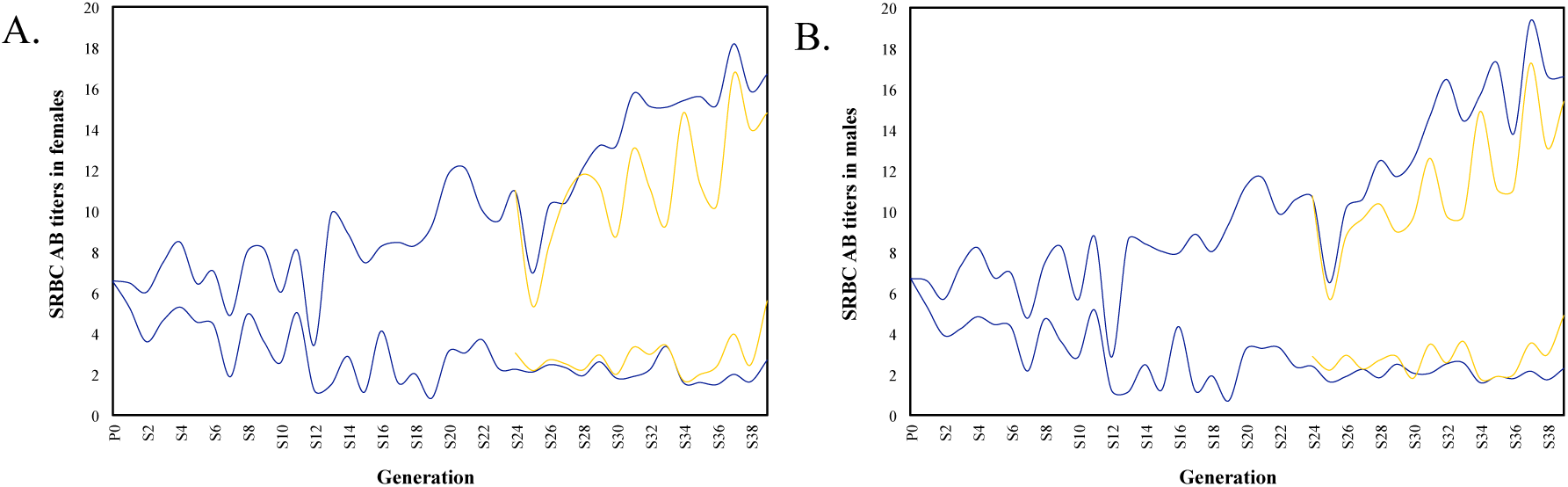
Changes in mean sheep red blood cell (SRBC) antibody (AB) titer across 39 generations of selection in Virginia AB chicken lines for females (A) and males (B). Selected lines shown in blue, with relaxed lines shown in yellow.

We observed hundreds of regions across the genome affected by the selection regime, as characterised by spans of extreme differentiation between HAS39 and LAS39. Despite the possibility of a strong confounding influence of drift in these relatively small experimental populations, the extent of genomic differentiation greatly exceeded predictions of genetic drift from population simulations, implying that many loci have undergone divergent selection and contributed to the selection response. To identify the most interesting candidate sweep regions, we overlap the sweep regions with associations detected to antibody response in an F_2_ intercross beween the selected lines, use enrichment analysis to detect sweeps with immune genes and identify sweeps located within genomic regions associated with immune traits in other populations. In this way, a high-confidence set of candidate selective sweep regions can be identified, and three particularly interesting candidate regions are presented in more detail.

## Materials and Methods

### Experimental populations: the long-term bi-directionally selected Virginia Antibody chicken lines

The Virginia Antibody chicken lines were established from the Cornell randombred White Leghorn population (Siegel and Gross 1980). From this base-population, two bi-directionally selected lines have been bred for high (HAS) and low (LAS) antibody response to an intravenous inoculation of 0.1 ml of 0.25% suspension SRBC, administered between 41 and 51 days of age. Plasma was collected five days after the inoculation and antibody response measured through a simple hemagglutination assay(Wegmann and Smithies 1966). Selection has been carried out once every year, with approximately 120 chicks hatched per generation. Between generations 1-10, 7 males and 28 females were selected to produce the next generation of each line, and from generation 11 onwards, 8 males and 32 females were used. Restricted truncation selection was used in each generation to reduce inbreeding (Kuehn et al. 2006; Zhao et al. 2012). Response to selection continues in HAS, whereas the LAS appears to have reached a selection-plateau (Figure 1). This plateau may be due to a threshold in response to the SRBC challenge or a limit in the technical sensitivity of the standard hemagglutination assay employed in antigen quantification (Dorshorst et al. 2011; Zhao et al. 2012). Relaxed sublines from both the HAS and LAS lines were established in generation 24.

The relaxed line from the high (HAR) and low (LAR) antibody response lines were founded by randomly selecting parents from the HAS and LAS lines, respectively.

DNA for the genomic analyses was prepared from blood samples collected from between 16-30 individuals from each line and pooled in equimolar ratios prior to library construction (Table 1).

**Table 1.** Information about the populations, samples, phenotypes and sequencing-depth used in the genomic analyses of the Virginia Antibody chicken lines.

| Population | N | Day 5 log<sub>2</sub>(AB titers) |  | Genome Coverage |
| --- | --- | --- | --- | --- |
|  |  | Mean | sd |  |
| HAS39 | 30 | 16.67 | 4.27 | 32.3 |
| LAS39 | 30 | 2.56 | 1.59 | 36.7 |
| HAR16 | 20 | 15.02 | 3.16 | 35.4 |
| LAR16 | 16 | 5.29 | 2.25 | 34.8 |

*Population: H-High, L-Low, A-Antibody, S-Selected, R-Relaxed, 39-generation; N: Number of individuals in the sequenced pool; Genome Coverage: average sequence-depth of the pool; AB: Antibody*.

### Pooled whole-genome re-sequencing, sequence alignment, variant-calling and population genomic analyses

Genome sequencing library construction and sequencing was carried out by SciLifeLab (Uppsala, Sweden) using two lanes on an Illumina Hiseq 2500. Reads were aligned to the *Gallus gallus* genome (Galgal4; INSDC Assembly GCA_000002315.2, Nov 2011) using BWA (Li and Durbin 2009). Picard (v1.92; http://picard.sourceforge.net) was used to sort genomes and to mark and remove duplicates. GATK (v3.3.0; McKenna et al. 2010) was used for realignment around indels. Samtools (v1.1; Li 2011; Li et al. 2009) was used to generate mpileup files for population genomic comparisons. GATK UnifiedGenotyper was used to generate allele calls at all sites (option: emit all sites) and with ploidy = 30 to account for the pooled genome sample. Sites were filtered to only include those with >10 and <100 reads, wherefrom allele frequency, heterozygosity, and pairwise *F*_*ST*_ between all populations were calculated. PoPoolation2 (v1.201; Kofler et al. 2011) was used to calculate *F*_*ST*_ over 1000 bp sliding windows with 50% overlaps between the population samples using the Karlsson et al. (2007) method, with minimum count 3, minimum coverage 10, maximum coverage 100, and minimum coverage fraction 1.

### Estimating the expected contribution of genetic drift to the genomic divergence between the populations

Our experimental populations are small (*Ne*_(1-10)_= 22.4 and *Ne*_(11-39)_= 25.6), and thus will increase the influence of genetic drift on shaping allele frequencies within populations and confound identification of selective sweep signatures. Therefore, simulations were used to estimate the expected contribution of drift to the genomic divergence between the lines. As input for these simulations, we used an estimate of the expected number of independently segregating regions in the final generation (39) in the HAS and LAS lines. This estimate was obtained by computing the expected population recombination rates in a random-mating 39 generation pedigree, under the assumption that drifting loci segregate independently. Based on this estimation, we expect that 1,408 regions in the genome segregate independently at the final generation. Then, 10,000 simulations of drift were conducted using the Wright-Fisher model on 1,408 randomly segregating bi-allelic loci, accounting for the changes in breeding-population sizes. Allele frequencies in the final simulated generation 39 were summarised and the differentiation due to drift computed. Two sets of starting allele frequencies were used: i) all loci at p = q = 0.5 to obtain an upperbound estimate for the effects of drift on the genome, and ii) randomly assigned starting allele frequencies from a uniform distribution to obtain a realistic estimate for selection starting from a random-bred population.

### Identification of candidate selective sweep regions

Regions demonstrating high population differentiation (*F*_*ST*_) between the HAS and LAS lines were considered as candidate selective sweeps. Based on the result of the drift simulations, a very stringent *F*_*ST*_ cutoff (> 95% percentile *F*_*ST*_ = 0.946) was used when defining the sweeps to limit the number of candidate regions due to drift. Windows with *F*_*ST*_ values above this cutoff were clustered into candidate sweep regions when they were less than 0.5 Mb from one another (custom R scripts). Clusters containing only a single 1000 bp window or less than 2 SNPs were removed from the candidate sweep dataset.

### Gene ontology analysis to identify immune genes within sweep regions

We used GO analysis to identify functional candidate selective sweeps that were enriched for immune-related genes. BEDOPS (Neph et al. 2012) was used to extract all Ensembl geneIDs underlying candidate sweep regions then used for GO annotation analysis via i) the Database for Annotation, Visualization and Integrated Discovery (DAVID v6.7; Huang et al. 2009) and ii) the Protein ANalysis THrough Evolutionary Relationships (PANTHER v10.0; Mi et al. 2016; Thomas et al. 2003). GeneIDs with immunological GO terms were identified and mapped back onto candidate sweep clusters.

To confirm transcription of the genes predicted by Ensembl genebuild within candidate sweep regions, White Leghorn spleen RNA sequence reads were accessed from the Short Read Archive (http://www.ncbi.nlm.nih.gov/sra; runs SRR2889287, SRR2889288, SRR2889289) and aligned using STAR (v2.3.1; Dobin et al. 2013). Genome coverage of the RNAseq alignments was calculated with BEDTools (v2.25.0; Quinlan and Hall 2010). RNAseq assemblies and genome coverages was visualised in IGV (v2.3.52; Robinson et al. 2011; Thorvaldsdottir et al. 2013).

### Identification of associations between candidate selective sweep regions and immune-traits

A number of QTL and GWAS studies have earlier been performed for immunological traits in chickens, including an analysis of an F_2_ intercross between the HAS and LAS populations. Here, we utilized the data and results from these studies in two ways to increase the confidence in individual candidate sweep regions. We reanalysed the SNP association data from Dorshorst et al. (2011) to compare the overlap between associations in those data and our candidate selective sweeps using a multi-locus, adaptive backward-elimination model-selection approach (Sheng et al 2015). Briefly, this dataset consisted of 128 individuals representing the phenotypic extremes in 5-day antibody titer sampled from an F_2_ intercross generated by intercrossing birds from HAS32 and LAS32 that were genotyped by a custom GoldenGate^®^ Genotyping assay containing 3,072 SNP markers as described by Dorshorst et al. (2011).

In order to focus on the most informative markers that tag divergent selective sweep regions, we used a subset of SNP markers that possessed an allele frequency difference > 0.7 in the pooled genome sequence between HAS39 and LAS39. To further refine this subset, neighbouring markers were clustered together (< 5 Mb between SNPs in a cluster, > 5 Mb between clusters). Backward elimination was applied to clusters with more than one marker to select the most significant marker as representative of the cluster region to retain in the genome-wide analysis. This final refined SNP subset was analysed using the same backward elimination process as in the clusters. This analysis uses a standard linear model framework, starting with a full model including the fixed effects of sex and the additive effects of the selected highly differentiated markers that were regressed on to log_2_ transformed day 5 antibody titers, and the final model was decided using an adaptive 5% FDR criterium (Abramovich et al. 2006; Gavrilov et al. 2009).

### Estimating haplotype segregation within candidate sweep regions of interest

This experimental population has a well-documented population history and extreme genomic differentiation within the defined candidate sweep regions. Appropriately, we are afforded the opportunity to disentangle highly divergent haplotypes based on allele frequency differences observed in the pooled genome sequencing data. Where both lines are fixed for different haplotypes, these can be identified by homozygosity throughout the region and haplotypes can be determined by adjusting all allele frequencies to the reference haplotype from one line. In other cases, one line is fixed within a sweep region, while different haplotypes continue to segregate in the other line. Here, positions are sequentially filtered on an ad hoc basis by allele frequencies differing from the selected reference hapslotype present in the fixed line. Allele frequencies in all lines are adjusted to reflect alternate allele frequencies from the reference haplotype, allowing visualisation and inference of alternative segregating haplotypes (see also Figure S1). As shown in the results, this approach was useful to estimate haplotype-frequencies at two candidate genes.

## Results

### Estimating the contribution by drift to the allelic divergence between populations

We used simulations to estimate the expected contribution by genetic drift to the genome-wide divergence between the divergently selected populations in the Virginia Antibody lines (Figure S2). When a uniform distribution of starting allele frequencies was assumed for the 1,408 regions that were estimated to segregate independently in the genome of HAS39 and LAS39, our similations (> 50,000 replicates) indicate that on average 47 loci (3.3% of all) are expected to become fixed for alternative alleles in the divergent lines due to drift, and that 53 loci would reach an *F*_*ST*_ > 0.946 (3.8%). When assuming a starting allele frequency of 0.5 for all 1,408 loci, which maximizes the risk of randomly divergent fixation of loci in the lines, on average 81 loci (5.7%) were fixed for alternative alleles. Here, we used *F*_*ST*_ > 0.946 (the 95% percentile of *F*_*ST*_ values between HAS39 and LAS39) as the cutoff for identifying candidate selective sweeps.

### A large genome-wide footprint of selection in the divergently selected lines

A total of 711,149 1,000 bp windows was analysed in *Popoolation2*. A summary of the population-statistics for the four analysed populations are provided in Table 2. Here we observed a general reduction in heterozygosity in sweeps relative to genome-wide heterozygosity in the selected lines. Differentiation (*F*_*ST*_) between lines was also greater between the S39 populations than R16 lines.

**Table 2.** Summary statistics on pooled genomes of Antibody Line populations

|  | HAS39 | LAS39 | HAR16 | LAR16 |
| --- | --- | --- | --- | --- |
| Heterozygosity in Sweeps | 0.120 | 0.127 | 0.191 | 0.156 |
| Genome-wide | 0.199 | 0.190 | 0.204 | 0.166 |

| Heterozygosity |  |  |  |  |
| --- | --- | --- | --- | --- |
| Sweep Het/ Genome Het | 0.603 | 0.666 | 0.939 | 0.940 |
| $F_{ST}$ | 0.519 | | 0.323 | |

After clustering of windows located less than 0.5 Mb apart and removing sweep-regions with a single 1,000 bp window or only 2 SNPs, 224 candidate sweep regions were retained (Figure 2; Table S1). These sweeps were located on 50 genome contigs, with 203 sweep regions across the 29 assembled chromosomes and 1 region on each of 21 unmapped genome scaffold, covering a total of 208.8 Mb (20.1% of the assembled galgal4 chicken genome). The regions ranged in length from 1.5 kb to 8.7 Mb (mean/median length: 932/538 kb).

**Figure 2.**
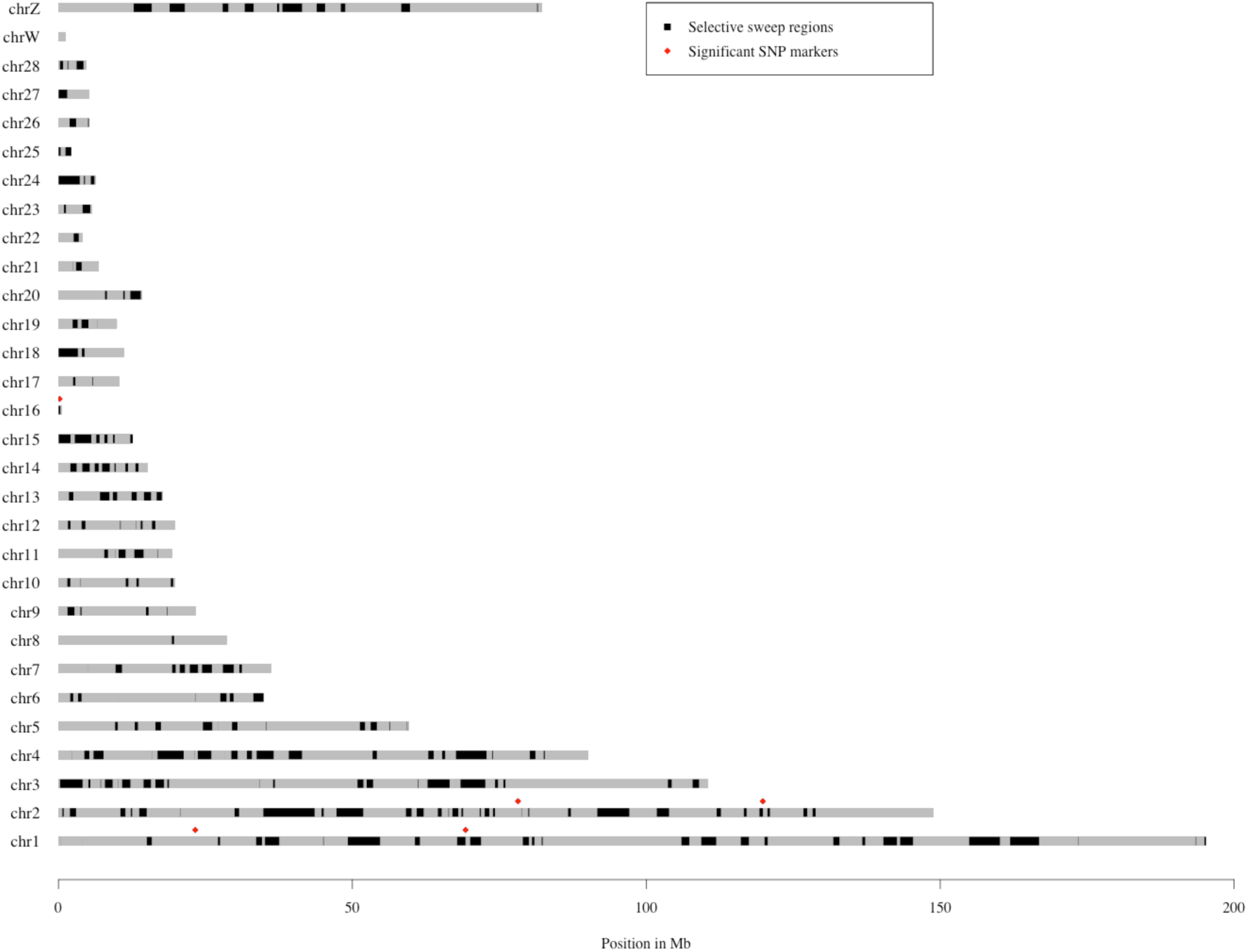
Distribution of candidate sweep regions (black) across chicken chromosomes. The selective sweeps associated with 5-day antibody titer in our backward-elimination based analysis of the *F*2 intercross between HAS32 and LAS32 is indicated with red diamonds.

### Many candidate selective sweep regions contain genes related with immune-function

The 224 candidate selective-sweeps covered in total 3,511 annotated genes. To identify potential candidate adaptive genes, as well as to indicate individual candidate sweeps that are more likely to have reached high frequency by selection rather than drift, we subjected these to a Gene Ontology (GO) analysis. Using the DAVID GO analysis, 67 gene IDs were reported to be associated with immune system processes (Table S2), and these mapped to 46 candidate sweep regions (Table S1). The PANTHER GO analysis identified 155 gene IDs associated with immune function (Table S3) mapping to 70 candidate sweep regions (Table S1). When combining the results from both methods, a total of 82 candidate sweep regions were found to include genes associated with immune function. Between 1 and 13 immune genes were found within these regions, with the most observed on GGA 16, notably the MHC region.

### Several candidate selective-sweep regions overlap with genomic regions associated with immunological traits in chicken

In total, 23 candidate sweep regions overlapped with genome regions that have been reported to be associated with immunological traits in chickens, including antibody response to SRBC and *Brucella abortus* in Leghorn hens (Zhou et al. 2003), primary and secondary antibody response to SRBC in ISA Warren layer hens (Siwek et al. 2003), innate immunity in layer hens (Siwek et al. 2006), innate and adaptive immunity in layer hens (Biscarini et al. 2010), multiple immune traits in the Chinese indigenous breed Bejing-You chicken (Zhang et al. 2015), and differential expression between high and low SRBC antibody responses in White Leghorn females (Geng et al. 2015) (Table S1).

### Several of the candidate selective-sweep regions are associated with day 5 antibody titers in an F_2_ intercross between chickens from HAS32 and LAS32

We reanalyzed a previously generated dataset from an F_2_ intercross from generation 32 of the divergent lines. In total, 150 of the 1024 polymorphic markers were highly differentiated, with an allele-frequency difference between HAS32 and LAS32 > 0.7. This SNP subset was clustered into 63 regions spanned 24 chromosomes. This subset of 63 representative SNP markers was fitted jointly in a whole-genome in a multi-locus, backward elimination analysis to identify five SNP markers associated with the selected trait at 20% FDR threshold, while four of these markers were retained in the model at the 5% FDR threshold (Table 3). These markers are located on chromosomes 1, 2, and 16.

**Table 3.** Genetic effects of SNP markers associated with log_2_(Day 5 Antibody titers) at a 5% FDR threshold in the F_2_ intercross of Virginia Antibody lines.

| *Marker* | *Chromosome<sup>a</sup>* | *Position<sup>b</sup> (bp)* | *Estimate<sup>c</sup>*<br>*± std. err.* | *P-value<sup>d</sup>* | *Sign.<sup>e</sup>* |
| --- | --- | --- | --- | --- | --- |
| rs14799859 | GGA 1 | 23,275,033 | -1.73 ± 0.54 | 0.0017 | 5% |
| rs15307852 | GGA 1 | 69,259,824 | 1.46 ± 0.46 | 0.0019 | 5% |
| rs14207559 | GGA 2 | 78,180,370 | -2.07 ± 0.43 | 3.6 x 10<sup>-6</sup> | 5% |
| rs14242328 | GGA 2 | 119,824,660 | 1.56 ± 0.44 | 0.0006 | 5% |
| rs14096690 | GGA 16 | 192,620 | -1.21 ± 0.42 | 0.0050 | 20% |

<sup>a</sup>GGA: Gallus gallus autosome
<sup>b</sup>November 2011 (galGal4) assembly
<sup>c</sup>Additive genetic effect in model including all tabulated loci

^a^ GGA: Gallus gallus autosome

^b^ November 2011 (galGal4) assembly

^c^ Additive genetic effect in model including all tabulated loci

^d^ Significance for additive genetic effect in model including all tabulated loci

^e^ Significance thresholds 5/20 % FDR based on which markers were selected

The marker with the most significant association, rs14207559, is located close to the sweep region at GGA2: 78,748,000 – 78,800,500, which contains the candiadate immune gene, semaphorin 5A *SEMA5A* (see below section). Among the markers selected by the 5% FDR threshold, rs14799859 is located on GGA1 between a sweep region ended at 15,887,000 and another beginning at 27,122,500, marker rs15307852 is located just outside the sweep region at GGA1: 67,820, 500 – 69,251,000 and marker rs14242328 falls within the sweep region on GGA2: 119,248,000 - 119,841,000, none of which contain candidate immune genes. The marker rs14096690 showed only 20% FDR significance in the model, but is located the sweep region GGA16: 2,000 – 323,000 and notably within the candidate Major Histocompatibility Complex region (see below section).

### SEMA5A is a candidate adaptive gene in a selective sweep region associated with day 5 antibody titer

A SNP marker rs14207559 that is significantly associated with day 5 antibody titer is located only 568 kb away from the candidate sweep region GGA2: 78,748,000 – 78,800,500 (Figure S3). It is fixed for T in HAS39, while it segregates for T/G in LAS at differentiated frequencies (0.27/0.73: T/G, Table 4). We explored the haplotype-structure of this selective-sweep in the four analysed populations in more detail where the sweep region overlaps with the *SEMA5A* candidate gene (GGA2: 78,760,000-78,800,000 bp; Figure 3). In this region, 345 positions are fixed in HAS39, but continue to segregate in LAS39 (Figure S3A/D). These positions in LAS39 contribute towards two major haplotypes, *Haplo2* and *Haplo3*, which are not present in HAS39, and segregate at the approximate haplotype frequencies of 0.2 and 0.8, respectively (Table 5, Figure 3). When compared to the relaxed lines, both HAR16 and LAR16 have *Haplo1* and *Haplo2* segregating at intermediate frequenices, and lack the *Haplo3* haplotype (Table 5, Figure 3).

**Table 4.** Estimates of mean phenotypes for the three genotypes at the significantly associated SNP near SEMA5A (rs14207559; GGA2: 78,180,370) in the F_2_ intercross between HAS32 and LAS32.

| *Genotype* |  |  |
| --- | --- | --- |
| *(rs14207559)* | *Mean antibody titer* | *SE* |
| G/G | 2.20 | 0.70 |
| G/T | 4.79 | 0.54 |
| T/T | 6.52 | 0.54 |

**Table 5.**
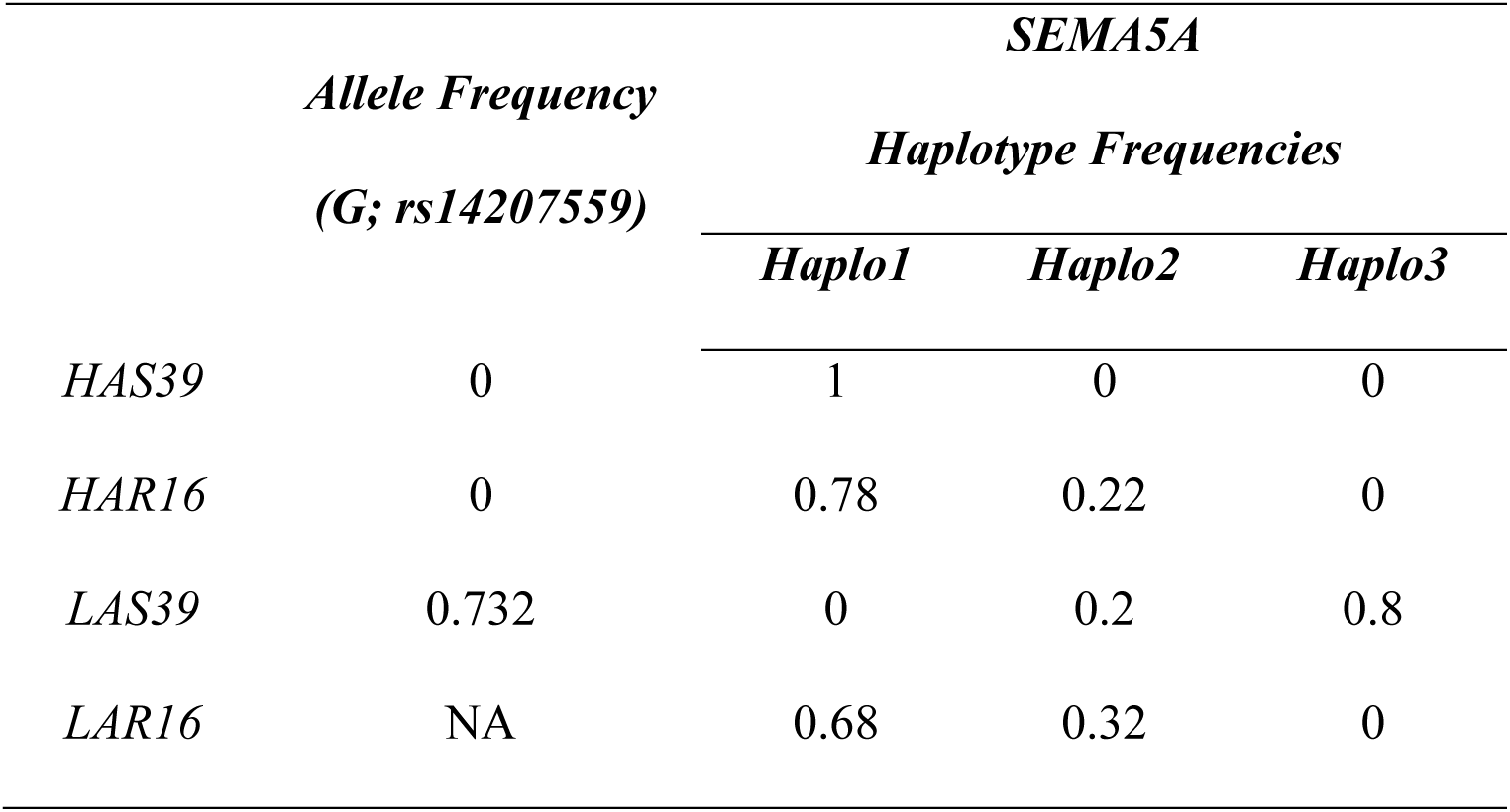
Allele and haplotype frequencies of the SNP marker and the SEMA5A gene-region.

**Figure 3:**
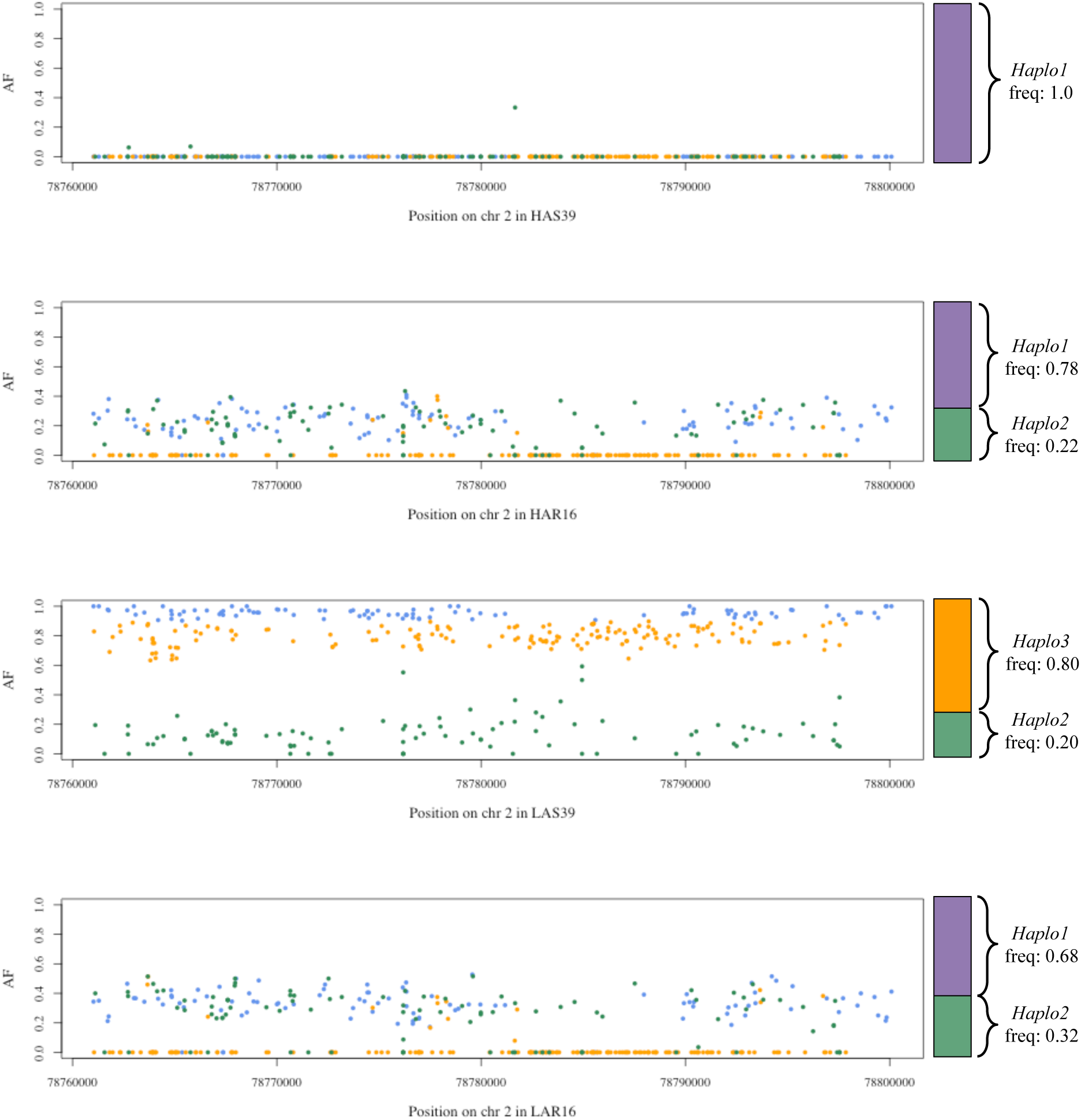
Inferred major SEMA5A haplotypes in HAS39 (upper), HAS 16 (middle upper), LAS39 (middle lower), LAR16 (lower). Positions are coloured based on their contibution to different haplotypes: allele frequencies in green contribute to *Haplo2*, orange contibute to *Haplo3*, blue contribute to both *Haplo2* and *Haplo3*, so where both of these haplotypes are present, blue positions will have allele frequency approximately equal to 1 as in LAS39; where only one haplotype is present, in the case of LAR16 and HAR16, the blue positions will have allele frequency approximate to the *Haplo2*.

The associated SNP marker rs14207559 appears to tag these major haplotypes at this selective-sweep region in the F_2_ population, with the G-allele tagging *Haplo3*, the dominant haplotype in LAS39, and the T-allele tagging both *Haplo1* and *Haplo2* (Table 5, Figure 3). From these inferred haplotype frequencies, it appears as though *Haplo3* possesses a selective benefit only in LAS and would otherwise be deleterious, which was purged in the LAR line after the relaxation of selection. Pairwise differentation of this region reinforces this hypothesis, as LAS39 is highly differentiated from HAS39, HAR16, and LAR16, while LAR16 appears less differentiated from HAS39 and HAR16 than LAS39 (Table 6). The several sequence variants within SEMA5A exons are synonymous, with the majority of polymorphisms in this region located in intergenic and intronic regions.

**Table 6.** Pairwise differentiation (mean F_ST_) between the analysed lineages across the region overlapping the SEMA5A gene and the candidate selective sweep (GGA2: 78,760,824 −78,800,500)

|  | *HAS39* | *LAS39* | *HAR16* | *LAR16* |
| --- | --- | --- | --- | --- |
| *HAS39* |  |  |  |  |
| *LAS39* | 0.595 |  |  |  |
| *HAR16* | 0.137 | 0.456 |  |  |
| *LAR16* | 0.202 | 0.438 | 0.029 |  |

### Longest sweep region overlaps with a deletion in *TGFBR2*

Our earlier work showed a correlation between the length of selective sweeps and their contribution to adaptation in a similar long-term, single-trait selection-experiments (Sheng et al. 2015). We therefore investigated the longest candidate sweep region (GGA2: 34,856,000 - 43,580,000; 8.7 Mb) in more detail. In this region, HAS39 and LAS39 are fixed for alternative haplotypes across this region, as are the respective relaxed lines. This pattern of fixation indicates that this region was fixed for alternative alleles within the selected lines prior to generation 24, which was the founding generation of the relaxed lines. Such rapid fixation suggests that this sweep might represent selection on a standing variant with strong phenotypic effect. Several genes with roles in the immune system are found within this region, including several cytokines, and we also detected a large structural variant within the transforming growth factor beta receptor 2 gene, *TGFBR2* (ENSGALG00000011442, GGA2: 40,385,525 - 40,447,574).

By referring to RNA seq data, we identified 9 exons contributing to the transcription of *TGFBR2* (Figure S4). Exons 2-3 and 4-5 of *TGFBR2* appear to be the result from a duplication. Exons 3 and 5 are identical at the sequence level, while exons 2 and 4 share over 90% sequence identity (intron 2-3 shares > 94% sequence identity with intron 4-5). The haplotype fixed in LAS39 to have a 3 kb deletion overlapping exons 4-5 of this duplicated region (GGA2: 40,414,500 - 40,417,500; Figure S4). This haplotype is also fixed in LAR16, implying that this deletion haplotype was fixed or close to being fixed by generation 24.

### Major Histocompatibility Complex

Several studies have previously investigated the difference in MHC allele segregation between the HAS and LAS lines. As such, we simply aimed to add information from the pooled genome sequencing to these results. We confirmed fixation for MHC *B* locus *B*^*21*^ haplotype in HAS39 and *B*^*13*^ in LAS39 (Table 6). We also observed fixation for *B*^*13*^ in LAR16 but interestingly HAR16 continued to contain both *B*^*21*^ and *B*^*13*^ haplotypes segregating at the approximate haplotype frequencies 0.73 and 0.27, respectively (Table 6).

**Table 6.** MHC B locus B^21^ and B^13^ haplotype frequencies for different generations in the Virginia Antibody lines. Note that allele frequencies in HAS10 do not equal 1, as a third MHC haplotype B^31^ was present at a frequency of 0.05 in this generation (Martins et al 1990).

| Generation | HA |  | LA |  | Citation |
| --- | --- | --- | --- | --- | --- |
| | $freq(B^{21})$ | $freq(B^{13})$ | $freq(B^{21})$ | $freq(B^{13})$ | |
| S10 | 0.80 | 0.15 | 0.01 | 0.99 | Martin et al 1990 |
| S13 | 0.99 | 0.01 | 0.02 | 0.98 | Martin et al 1990 |
| S32 | 1 | 0 | 0 | 1 | Dorshorst et al 2010 |
| R16 | 0.73 | 0.27 | 0 | 1 | This study |

## Discussion

Long-term, bi-directional experimental selection is a powerful approach to gain empirical insights to the genetics of complex traits. Within the quantitative genetic framework, research has provided knowledge about relationships between the predicted and realized responses to selection, the boundaries of selection in closed populations due to reduction in selectable genetic variation and physiological limits for further response (Hill 2010; Dunnington and Siegel 1996). With the advent of large-scale molecular genetics and genomics methods, these highly selected populations also become valuable models for studying how the genomes of populations under intense selection on various traits evolve during adaptation (Tobler et al. 2014; Barrick et al. 2009; Burke et al. 2010; Johansson et al. 2010)

The Virginia antibody lines are a unique experimental model-population for studying the genetics of long-term selection for an adaptive immune trait in a higher vertebrate. Using whole-genome re-sequencing of pooled DNA from these divergent lines, we revealed wide genome involvement in the long-term selected response. Over 200 regions, covering nearly 20% of the genome, were clearly differentiated between the lines. The presence of such extensive selective signatures in the genome is primarily the result of two key factors: firstly, the selected variants were present at the onset of selection (i.e. that selection has acted on standing genetic variation in the founding population), and secondly that selection acts on these standing variants in opposite directions. Even in smaller breeding populations, where random genetic drift will play a large role, using simulations we estimate that most (about 81%) of the observed genomic differentiation between the lines is expected to be due to selection, assuming that the allele-frequencies at the selected loci were uniformly distributed in the base population.

These extensive genome changes in this long-term selection experiment thus strongly suggest that SRBC antibody response is a highly polygenic trait. The 6.5 fold difference in antibody titers between the divergently selected lines has been facilitated by large differentiation at many loci with standing genetic variants. This finding is consistent with the findings in the Virginia body weight lines, which has been bi-directionally selected for juvenile body-weight for over 55 generations. There it has been shown that selection on standing variation in a highly polygenic genetic architecture was the main contributor to the long-term selection response in body weight (Sheng et al. 2015).

Genetic drift in our small breeding populations would have contributed not only to random divergence between the selected lines, but also to the fixation of selective sweeps. Previous studies where *Drosophila melanogaster* and *Saccharomyces cerevisiae* were used to model vertebrate populations have rarely detected complete selective sweeps (Burke 2012; Burke et al. 2014). Further, in human studies, the polygenic architecture of most traits disperses the influence of selection and hence adaptation is faciltated by moderate allele frequency changes of multiple loci without dramatic fixation events (Coop et al. 2009; Pritchard et al. 2010). In contrast, within our small population, there was loss of heterozygosity within selective sweep regions attributable to fixation of genetic variants. The combination of selection and genetic drift within the Antibody lines have lead to a higher level of fixation of selective sweeps than observed in other model systems.

Antibody production is a major defence mechanism of the humoral immune response. It involves a variety of different cell types with specific functions, including immunological cascades involving the recognition of the foreign cells by T and B immune cells, activation and differentiation of B cells into plasma cells and secretion of specific antibodies (Janeway et al. 2001). Moreover, this cascade relies on the careful coordination of gene expression through signalling molecules and appropriate receptors, within which there is potential for many genes and genetic variants to have an effect. The identification of a large number of genomic regions that are fixed for alternative allelic variants in these lines suggest not just that many loci play a role in SRBC antibody production, and also that the founder population harboured substantial genetic variation upon which selection could act. This supports the expectation that adaptive immune traits are genetically complex, and further emphasises the need to consider more than just a few key immune genes when studying immunogenetic selection and adaptation even in breeding populations originating from a narrow genetic basis.

Earlier work in the Virginia Antibody lines have, however, shown that the level of antibody-response does not have a direct connection to a general greater immunological efficacy. For example, when the lines selected for high (HAS) and low (LAS) antibody production were faced with different immunological challenges, the HAS were less susceptible to the northern fowl mite (*Ornithonyssus sylviarum*) and Newcastle Disease, but more susceptible to the bacterial pathogens *Escherichia coli* and *Staphylococcus aureus* when compared to LAS (Gross et al. 1980). These immunological trade-offs further emphasis the complexity of the immune system, and show fixation of mutations within this selected population that would be otherwise deleterious. Further, they also serve as a reminder that this study does not provide insights to how and whether breeding can be used to improve the general performance of the immune system, but rather that the immune-system of a particular population is likely to be very adaptible when challenged with various disease challenges and that such responses are likely due to selection on available variants in many genes. It is advantageous for a population to have such variation in host-pathogen interactions.

We observed many genes with immunological function underlying our candidate sweep regions, while many still did not. Particularly from our association analysis, while we found two SNPs markers with strong association to day 5 antibody titers and closely located candidate immune genes, the remaining three SNP markers were not located near immune genes involved in selective sweeps. This may be a consequence of our strict GO search for genes with immune function, which would not identify functional genes contributing to the divergent antibody response but without GO immune function terms. In the remaining sections, we will discuss a number of particularly promising candidate functional genes detected in the identified sweep regions.

### Semaphorin-5A

A sweep region located near the SNP marker (rs14207559) with the strongest association to day 5 antibody titers in the F_2_ intercross between the HAS and LAS lines in generation 32 overlaps the first 7 of 18 exons (39,676 of 136,181 bp) of Semaphorin-5A (*SEMA5A*, ENSGALG00000013051). Semaphorins are a large family of secreted and membrane bound proteins, originally discovered in the nervous system with a role in axon guidance (Lyu et al. 2015). *SEMA5A* is a secreted semaphorin involved in immune regulation and the pathogenesis of autoimmune diseases (Lyu et al. 2015). It has been implicated in the pathogenesis of immunological diseases such as mastitis in dairy cattle (Sugimoto et al. 2006), chronic immune thrombocotypenia in humans (Lyu et al. 2015), and neurodevelopmental disorders such as autism in humans (Melin et al. 2006). As an immunoregulator, *SEMA5A* influences the expression of cytokines such as IFN-gamma, TNF-gamma and interleukins (Lyu et al. 2015).

In the Virginia Antibody lines, the sweep region overlapping the *SEMA5A* gene is present in at least three different haplotypes: the major haplotypes referred to here as *Haplo1*, *Haplo2*, and *Haplo3. Haplo3* is present only in LAS39, suggesting that the variant leading to low antibody production was purged in relaxed line LAR16. This is reflected in the population differentiation between the lines as well, as LAS39 is most differentiated of the lines. This extent of differentiation at many markers suggests that these haplotypes were present as standing variants in the base population, as opposed to a novel variant emerging in the LAS lineage after generation 24 and the founding of the relaxed line. We observed only three synonymous SNP sites in the coding region of *SEMA5A*, with the majority of sequence polymorphism between haplotypes contained in intergenic and intronic regions. Taken together this may suggest that the involvement of *SEMA5A* to the differential antibody response in the Virginia Antibody lines would be due to regulatory polymorphisms where the two haplotypes produce different expression levels of the *SEMA5A* protein, from different promoter or enhancer sequences, which could impact the expression of cytokines within this regulatory network, influencing an effective and specific immune response.

### Transforming growth factor beta receptor 2

Earlier work in the Virginia body-weight selected lines, which have undergone a similar long-553 term experimental selection regime, has shown that the length of a selective-sweep is correlated with its contribution to the adaptative phenotype. Hence, the individual sweep in our study that is most likely to make a significant contribution to antibody response is the long 8.7 Mb sweep region identified on GGA2. In this region, only one gene had GO immune terms, the transforming growth factor beta receptor 2 (*TGFBR2*), making it an interesting candidate gene for further study. *TGFBR2* is a member of the serine/threonine protein kinase family, encoding a transmembrane protein that binds the secreted protein TGF-beta. The TGF signalling pathway regulates diverse biological processes during all stages of life and plays a key role in growth and the immune response (Letterio and Roberts 1998; Li et al. 2006).

We observed fixation for different *TGFBR2* haplotypes in the selected lines. In-depth analyses comparing the sequence data for HAS39 and LAS39 suggests that these haplotypes differ by an approximate 3 kb deletion (between GGA2: 40,414,500 - 40,417,500) fixed in LAS39, which encompassed exons 4 and 5 of the *TGFBR2* gene prediction (Figure S4). However, the precise length of the deletion is difficult to resolve by the short reads of our pooled genome sequencing, and confounded by the historical duplication and deletion. Further analyses of the relaxed lines (HAR16 and LAR16) show that the HAS and LAS haplotypes were fixed for the respective variants also in the relaxed lines. This suggests that the lines were fixed (or nearly fixed) already by selected generation 24 when the relaxed lines were founded. The length of this sweep, together with the rapid fixation, suggests that the selection acted strongly and fixation occurred swiftly, before multiple recombination events could break down the linkage within this region and leave a shorter sweep.

The strong selection signature suggests that the *TGFBR2* haplotypes might have played a major role in determining antibody response, most likely due to its key function in the TGF-beta pathway. However, there was no evidence that this large sweep region was associated with 5 day antibody titers in the F_2_ intercross between HAS32 and LAS32, despite it being covered by 7 segregating SNP markers that were expected to be highly informative for the line difference in this region (6 markers with MAF > 0.05; 605,124 bp or greater distance from the *TGFBR2* annotated gene). It should be noted, however, that these SNP markers were fixed or near fixed for alternative alleles in generation 39. As the population used for association analysis was small, the power in this analysis was low, making it difficult to make strong conclusions as to whether these haplotypes had no or only minor effects on antibody response, or whether their functional significance was reliant on the genetic background of the population.

Defects in *TGFBR2* causes serious debilitating and fatal diseases in human, most notably the autosomal dominant aortic aneurysm syndrome, Loeys-Dietz syndrome, but also increased risk of cancers (Loeys et al. 2006; Levy and Hill 2006). That none of these conditions have been observed in LAS suggests that the deletion observed in LAS39 and LAR16 does not completely disrupt the function of the gene or that the function of this gene is less important in chickens than in humans. The deletion of only one set of duplicated exons may indicate that the LAS haplotype may still result in a functional *TGFBR2* protein, but with less splice variability than the HAS haplotype.

### Major Histocompatibility Complex Region

The major histocompatibility complex (MHC) is the most well studied immune gene complex in vertebrates due to its key role in the pathogen surveillance (Klein 1986), and accordingly the MHC has previously been investigated in the Virginia Antibody lines, albeit not at a sequence level (Dunnington et al. 1989; Martin et al. 1990; Dorshorst et al. 2011). The chicken MHC region is relatively small, simple and tightly-linked, compared to that of mammals, and thus termed the minimal essential MHC (Kaufman 2015; Kaufman et al. 1999). Bi-directional selection on the MHC was evident as early as generation 12 in the Virginia Antibody lines, and alternative haplotypes were fixed by generation 32 (haplotype frequencies are summarised in Table 6; Dorshorst et al. 2011; Dunnington et al. 1989). Fixation for *B*^*21*^ in HAS39 and *B*^*13*^ in LAS39 was confirmed in our pooled genome sequencing and this divergent fixation contributed to the sweep signal on GGA16 from 2,000 - 323,000 bp.

The sweep overlapping the MHC region spans a total 321kb, and encompasses genes vital to antigen processing and presentation including MHC class I and II, TAP genes and tapasin. Association analysis between MHC haplotype and antibody response has demonstrated that the *B*^*21*^ allele acts dominantly, with higher day 5 and day 12 titers of antibodies in homozygous and heterozygous *B*^*21*^ individuals (Dorshorst et al. 2011). This explains the rapid fixation for the recessive *B*^*13*^ in the LAS, as inferred by the fixation in LAR16, and why both haplotypes still segregate in HAR16. Interestingly, although the HAS13 was close to fixation for *B*^*21*^, at least one *B*^*13*^ haplotype must have persisted until generation 24 the origin of the relaxed lines, as it continues to segregate in HAR16. Balancing selection, known to act on the MHC region in natural populations (Charbonnel and Pemberton 2005; Hedrick 1998), may have been important in the recovery of haplotype frequencies from near fixation in HAS13 to heterozygosity in 0.394 in HAR16.

## Conclusion

Adaptation to long-term, bi-directional selection on antibody response in this experimental chicken population has been facilitated by standing genetic variation across many regions of the genome. Although selective-sweep studies by themselves are unable to quantify the contribution by individual genes and polymorphisms to adaptation, we use data from earlier association and functional studies to highlight three particularly interesting candidate sweeps and underlying candidate genes involved in immune surveillance (MHC) and immune regulation (*TGFBR2* and *SEMA5A*). Further work to investigate the large numbers of additional immune genes that are likely to have contributed to the response to selection in this experimental population would be useful to improve our understanding of selection on immune traits and to unravel the complexity of their interactions. These experimental lines continue to present a unique opportunity towards understanding the mode and tempo of selection in a higher vertebrate organism.

## Acknowledgements

Sequencing was performed by the SNP&SEQ Technology Platform, Science for Life Laboratory at Uppsala University, a national infrastructure supported by the Swedish Research Council (VR-RFI) and the Knut and Alice Wallenberg Foundation. Computations were performed on resources provided by SNIC through Uppsala Multidisciplinary Center for Advanced Computational Science (UPPMAX, project id: b2015010).

